# Impacts of white-tailed deer on regional patterns of forest tree recruitment

**DOI:** 10.1101/052357

**Authors:** Lauren Bradshaw, Donald M. Waller

## Abstract

Local, short-to medium-term studies make clear that white-tailed deer can greatly suppress tree growth and survival in palatable tree species. To assess how deer have broadly affected patterns of tree recruitment across northern Wisconsin, we analyzed recruitment success in 11 common trees species that vary in palatability across 13,105 USFS - FIA plots sampled between 1983 and 2013. We also examined how recruitment in these species covaried with estimated deer densities here. Saplings of five palatable species were scarce relative to less palatable species and showed highly skewed distributions. Scarcity and skew provide reliable signals of deer impacts even when deer have severely reduced recruitment and/or no reliable deer density data are available. Deer densities ranged from 2.3-23 deer per km^2^ over a 30 year period. Sapling numbers in two maples (*Acer*) and aspen (*Populus*) with intermediate palatability declined sharply in apparent response to higher deer density. Path analysis also reveals that deer act to cumulatively depress sapling recruitment in these species over successive decades. Together, these approaches show that deer have strongly depressed sapling recruitment in all taxa except *Abies* and *Picea*. As these impacts are now propagating into larger sized trees, deer are also altering canopy composition composition and dynamics. The tools developed here provide efficient and reliable indicators for monitoring deer impacts on forest tree recruitment using consistent data collected by public agencies.

## 1. Introduction

In the United States, forest products generate over $200 billion a year in sales nation wide (USDA Forest Service 2014). To maintain recreational and commercial use of these forests, managers must sustain forest growth by ensuring the responsible harvest of forest products and successful tree regeneration. Foresters often adjust management practices to enhance natural regeneration of desired species. Nevertheless, several ecological factors may act to inhibit seedling establishment, growth, and sapling recruitment. These factors include the often intense competition of seedlings and saplings for water, soil nutrients, and light (Aarssen & Epp 1990; Dalling et al. 2011). Thus, species traits like drought and shade tolerance strongly affect a tree’s ability to persist and compete for these resources. Seeds, seedlings, and saplings are also vulnerable to seed predation and a broad spectrum of herbivores including insects, birds, and mammals (Kolb, Ehrlen & Eriksson 2007). For some species like *Betula alleghaniensis*, a valuable timber species, the proportion of seeds that survive and persist into the sapling size classes is so low that these filters now serve to limit sapling recruitment and population persistence (Lorenzetti et al. 2008).

White-tailed deer (*Odocoileus virginianus*) now act as a key herbivore across much of northeastern North America limiting regeneration in many tree species (Rooney and Waller 2003; Côté et al. 2004). Deer consume seeds, seedlings, and the buds, flowers, leaves, and sometimes bark and branches of saplings in palatable woody species exerting strong impacts in winter and early spring. They prefer to graze on graminoids and palatable understory forbs in spring through summer (Healy 1971; Stormer and Bauer 1980; Berteaux 1998). Even when deer do not consume whole plants, their consumption of nutrient rich flowers, terminal meristems, and photosynthetic tissues tends to curtail growth and reproduction. Collectively and cumulatively, deer consume considerable understory biomass, strongly affecting energy and nutrient pathways. Ungulates also tread on plants and paw at leaf litter, destroying some plants and exposing mineral soil (Hobbs 1996; Persson et al. 2000; Russell et al. 2001). Deer can also act as seed predators and sometimes as vectors to disperse seeds (Ostfeld et al. 1996; Gill and Beardall 2001).

Through the latter 20^th^ century, populations of white-tailed deer increased across much of the Eastern and Midwestern United States. In Wisconsin, populations have increased several fold since the 1950s. Deer populations here now chronically exceed population goals in most Deer Management Units (WI DNR 2015a). Ecologists in the Upper Midwest (USA) first drew attention to the threats high deer populations pose to tree regeneration and forest plant communities in the 1940s and 1950s (Leopold et al. 1946; Dahlberg & Guettinger 1954). Later ecological studies confirmed the reality of these threats by demonstrating shifts in the abundance, height, and demographic profiles of species sensitive to deer herbivory (Anderson and Loucks 1979; Bratton 1979; Marquis 1981). In the Upper Midwest, the density and growth of several slow-growing, palatable woody species like *Tsuga canadensis*, *Betula alleghaniensis*, and *Thuja occidentalis* have declined sharply in regions with abundant deer (Rooney & Waller 2003; Rooney 2001; Waller & Alverson 1997). Understory forbs like *Trillium grandiflorum*, *Clintonia borealis*, and *Maianthemum canadense* also show strong declines where deer are abundant (Balgooyen & Waller 1995; Frerker et al. 2014).

Because trees take many years to mature, declines in long-lived trees often occur long after deer herbivory occurs. This means we may not witness its full impact for decades (McGarvey et al. 2013). Forests subjected to prolonged, elevated deer densities continue to reflect these impacts for 20-70 years after release from deer browse pressure (Nuttle et al. 2014; Anderson & Katz 1993; Balgooyen & Waller 1995). In addition, these shifts in plant community structure and composition also strongly affect the abundance of birds, small mammals, and other components of diversity (Allombert 2005; Cardinal 2012; deCalesta 1994; Fuller 2001; Martin et al. 2012; McShea and Rappole 2000; Ostfeld et al. 1996). These long-lasting impacts of deer herbivory could limit our ability to maintain and restore tree, understory plant, and animal diversity within North American forests. Such diversity supports significant ecological and economic values including populations of other wildlife species, an array of ecosystem services, recreational utility, and the timber value of commercially valuable hardwood species.

Ecologists use several methods to study deer impacts. These include tracking differences in growth rates, reproductive condition, the relative abundance of species, long-term shifts in community composition, natural experiments (e.g., islands with and without deer), and manipulative experiments (e.g., fenced exclosures) (Côté et al. 2004; Waller 2013). Although such studies provide valuable data, they are typically of short duration and apply primarily to the particular species that were studied and the local areas where the work was done. Exclosure studies that rigorously demonstrate strong deer effects on local plant communities are also sometimes criticized for making an extreme comparison between ambient deer effects and no deer at all. In addition, building and maintaining many exclosures over many years is expensive, often forcing us to rely on data from just a few exclosures at particular locations, reducing the generality of what we can infer. These concerns suggest that it would be useful to assess impacts of deer on tree regeneration across larger areas and a more natural range of variation in deer abundance. It would also be ideal if we could study deer impacts across multiple forest types, ages, and longer time periods, e.g., by linking local exclosure studies to longer-term regional trends (Frerker et al. 2014).

Here, we investigate regional variation in sapling abundance (recruitment) in 11 tree species in relation to variation in deer density that occurred over a 30 year interval and across all of northern Wisconsin. The broad scope of this study complements more intensive local short-term studies by providing a big picture of how deer are affecting tree recruitment in this region. To obtain this picture, we use systematic surveys of forest conditions pursued by the U.S. Forest Service in their Forest Inventory and Analysis (FIA) program (http://www.fia.fs.fed.us/). The FIA program surveys permanent plots arrayed on a regular grid at regular intervals (about 5-years). The number and dispersion of these plots (covering all forest lands in the U.S. since 1999) provide data of high statistical power from an unbiased set of samples. This allows us here to systematically compare variation in sapling recruitment among the chosen taxa in our region. In particular, we assess recruitment in 11 common tree species chosen to include taxa that differ in their palatability. These range from species that deer avoid (*Picea*) to species known to be highly palatable and susceptible to deer browsing impacts (*Thuja occidentalis* and *Tsuga canadensis*). We hypothesized that saplings of species that are more palatable and susceptible to deer would be: a) generally scarcer across the landscape, b) absent altogether from many sites, and c) scarcer at sites and in decades where they encountered higher deer densities. The deer density estimates we use for c) also derive from a public source, namely the Wisconsin Department of Natural Resources (Wis-DNR). Because the metrics and approaches we describe use only publicly available data systematically collected by professional agencies, they do not require forest or wildlife managers to acquire new data or conduct local research.

Many local factors also affect patterns of tree recruitment including local soil and light conditions, local canopy composition and seed inputs, local deer browse preferences, and tree harvest history. We lacked consistent data for these and also note that obtaining such data would be impractical for most managers. In addition, our goal here was to analyze variation in recruitment at the coarser spatial scale of whole DMUs in order to obtain reliable aggregated signals of deer impacts for making management decisions. Our coarser scale of analysis ensures such averaging by filtering out much of the “noise” generated by the many variable local factors also affecting tree recruitment and deer-tree interactions.

## 2. Methods

### 2.1 Study region and estimates of deer density

Our study region encompasses the northern third of Wisconsin dominated by mixed hardwood forestlands (Fig.1) providing a relatively homogenous set of landscapes for testing how deer affect tree recruitment. By limiting our study to this state, we could also use a single consistent set of deer density estimates from the Wis-DNR. They estimate annual post-hunt deer density using a Sex-Age-Kill (SAK) model in each of the 48 Deer Management Units (DMU) located in northern Wisconsin. Implementation of this model by the Wis-DNR provides relatively robust and reliable estimates of overwintering deer density with defined and limited amounts of error and bias (Millspaugh et. al 2009; T. VanDeelen, pers. commun.). Comparing SAK estimates with more rigorous Statistical Age-at-Harvest estimates that explicitly incorporate changing age structure and harvest rates indicates that the SAK model, as implemented in Wisconsin during our study period, tracked SAH estimates very closely in the northern forested DMUs (Norton et al. 2013). In addition, we only use the deer data in a comparative context to assess how variation in estimated deer numbers over DMUs and decades affects patterns of sapling recruitment in the chosen tree species.

**Figure 1.**
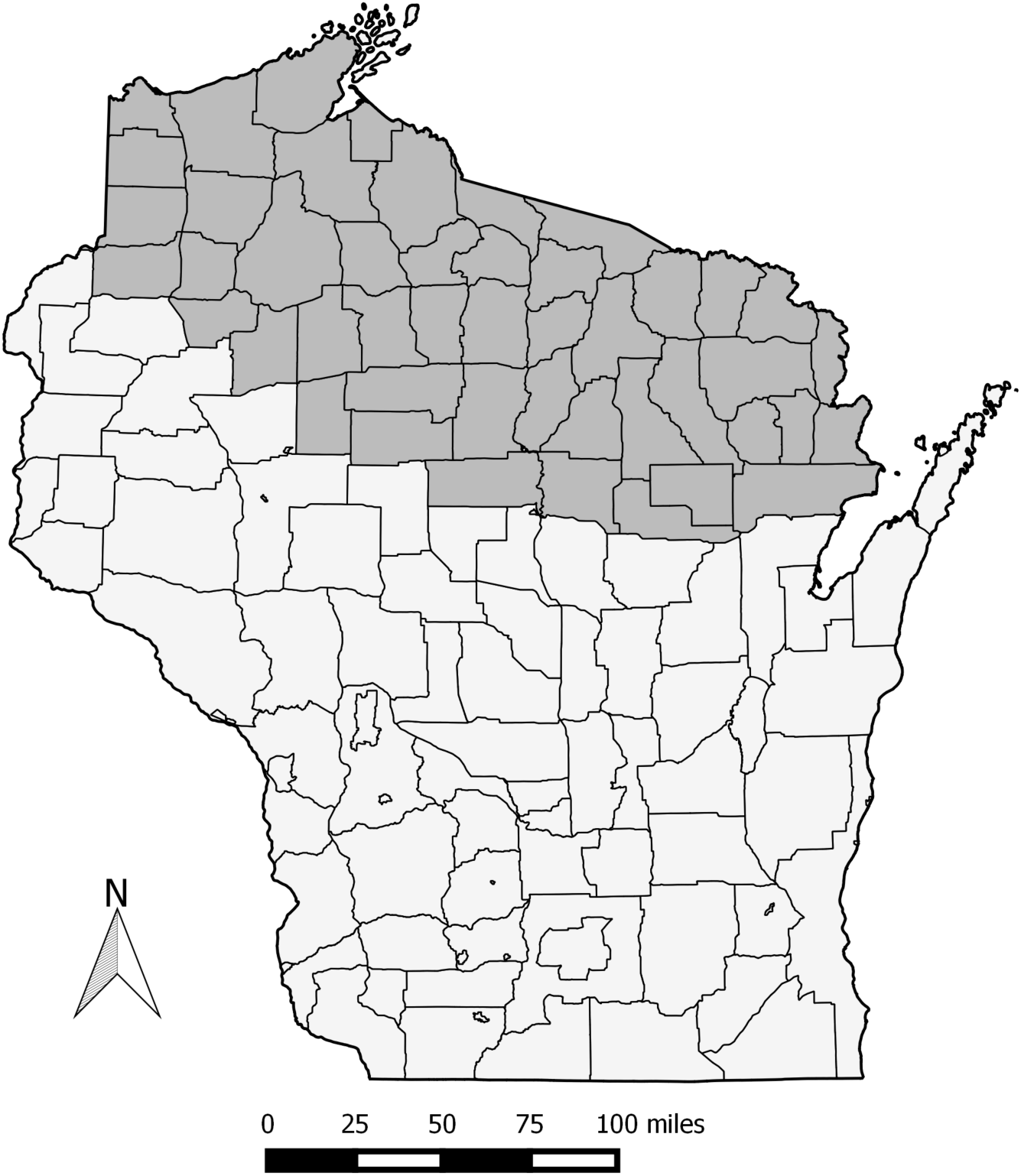
Study area (shaded) in northern mixed hardwood region of Wisconsin. Note that this area does not include the Apostle Islands.

The FIA plots occur at a density of about 1 plot per 1500 ha. We spatially divided these FIA plots into groups according to DMUs. The boundaries of these DMUs remained stable during the period of this study except for two large DMUs that were each split into two smaller units and one small DMU that was merged into a neighboring unit. For these, we recalculated estimated deer densities to match the new DMU areas. Our approach to hypothesis c) assumes that enough variation exits in deer density among the DMUs and study periods to alter the abundances of small saplings. Levels of variation in estimated deer density over these DMUs and the 30-year period are ample (Fig. 2), justifying the “natural experiment” approach we use here.

**Figure 2.**
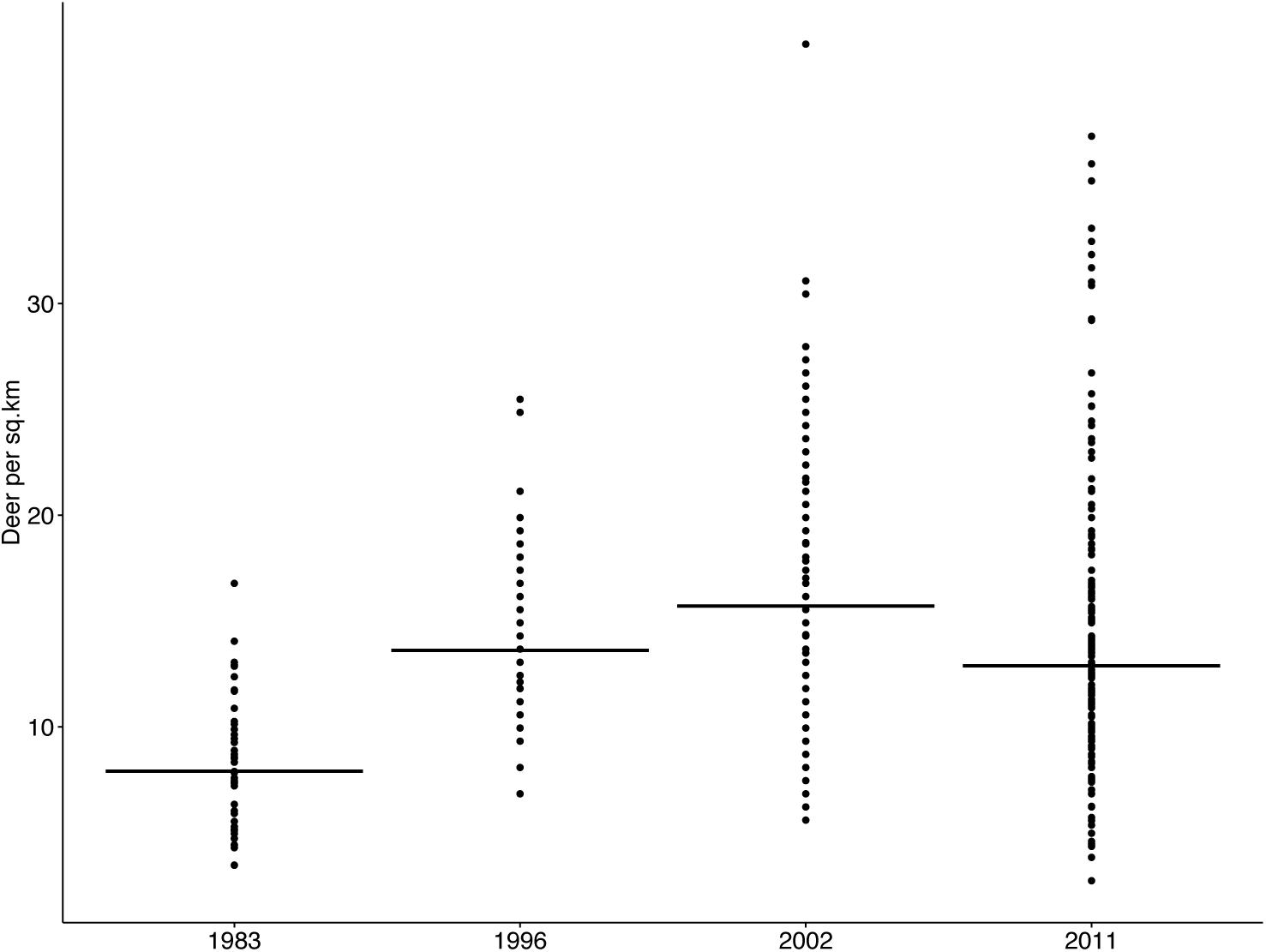
Variation in estimated deer densities (deer per km^2^) among Deer Management Units (DMUs) from 1983 to 2011. Note the high level of variation in estimated deer densities across DMUs and periods.

Studies typically assess the number or height of tree seedlings or the incidence of deer browsing on seedlings and saplings within the “molar zone” (generally taken to be 0-180cm above the ground). Such methods directly measure the immediate effects of browsing on local populations. Here, we sought instead to assess the cumulative impacts of prior deer browsing by instead tallying the number of small saplings (>2.54 cm DBH). The Phase 2 FIA data that we used ignores smaller saplings, making these the smallest trees for which we could obtain data. Trees >2.5cm DBH are well established, 3-5m tall, and have most of their foliage above the point where deer browse occurs (Kelty and Nyland 1981), making them far less vulnerable to deer impacts. These thresholds are often used in deer browse studies (White 2012; McGarvey et al. 2013, Bressette et al. 2012).

Because the number of FIA sampled saplings in a plot reflects a history of past browsing rather than current levels of browsing on smaller individuals in that stand, we analyzed variation in sapling recruitment in relation to estimates of deer density during the previous survey period (~10 years before the FIA sample year). The actual interval between when deer browsing occurs and the following FIA survey could be longer or shorter than this (likely longer in slow-growing species and perhaps shorter in fast-growing species). Given the decadal sampling regime and the broad scale of these surveys, we deemed 10 years to be the appropriate interval for evaluating deer effects. Analyses based on other lags showed weaker relationships to estimated deer density.

### 2.2 Study species and palatability

We focused on 11 common tree species known from previous studies to vary in their palatability to deer and sensitivity to browsing (Table 1). We specifically included two unpalatable genera (*Picea* and *Abies*) in order to provide a control that would allow us to detect any general spatial or temporal trend in tree recruitment unrelated to deer herbivory. We assigned the taxa to one of four palatability classes *a priori* based on published preference results for our region (e.g., Dahlberg and Guttinger 1956), a local exclosure study (Frerker et al. 2014), and our own expert opinion (based on 25 years of field work on deer impacts in Wisconsin and wide reading of the relevant literature). As *P. glauca* and *P. mariana* are similarly distasteful to deer and difficult to distinguish, we combined these into the genus *Picea*.

**Table 1.** Assigned browse sensitivity classes for the 11 focal taxa. Class 1 includes the least palatable species and Class 4 those that are most palatable and vulnerable to deer. Note the far higher skew in the abundance distributions of the (log) number of small saplings in the more palatable taxa relative to less palatable taxa or larger trees of the same species.

| Scientific Name | Common Name | Palatability<br>Class | Skew in log (# of<br>small saplings +1) | Skew in log<br>(# of large<br>trees +1) |
| --- | --- | --- | --- | --- |
| *Abies balsamea* | Balsam fir | 1 | 0.02 | 0.0283 |
| *Picea spp.* | Spruce | 1 | 0.60 | -0.1476 |
| *Acer rubrum* | Red maple | 2 | 0.82 | -0.501 |
| *Acer saccharum* | Sugar maple | 2 | 0.02 | -0.41 |
| *Populus tremuloides* | Quaking aspen | 2 | 0.04 | 0.3181 |
| *Pinus strobus* | White pine | 3 | **2.18** | -0.2047 |
| *Quercus rubra* | Northern red oak | 3 | **1.66** | 0.1289 |
| *Betula alleghaniensis* | Yellow birch | 4 | **1.47** | -0.16253 |
| *Thuja occidentalis* | Northern white<br>cedar | 4 | **2.52** | -0.1387 |
| *Tsuga canadensis* | Eastern hemlock | 4 | **2.00** | 0.25734 |

### 2.3 Assessing variation in tree recruitment

To obtain data on the abundance and sizes of trees for these ten taxa, we accessed the USDA National Forest Service Forest Inventory Analysis (FIA) “DataMart” via www.fia.fs.fed.us/tools-data. The FIA program resurveys long-term forest plots nation-wide. In Wisconsin, these commenced with intensive periodic surveys in 1983 and 1996 (roughly 1 plot per 1000-1500 ha). They then transitioned to less-intensive (roughly 1 plot per 7000 ha) annual surveys in 2000. We use tree data from 13,105 FIA plots in our region spanning the period 1983 to 2013 divided into four periods. To account for differences in sampling intensity across inventories, surveys from 2000-2004 and 2009-2013 were aggregated together to create similar plot densities (without duplication as plots are resurveyed on a 5-year cycle). We refer to these aggregations as inventory year ‘2002’ and ‘2011’, respectively. We divide trees in the FIA surveys into three size classes: small (2.54–5.08cm DBH), medium (5.08-10.16cm DBH), and large (>10.16cm DBH).

### 2.4 Statistical analyses

To address hypotheses a) and b), we first tallied numbers of saplings and trees in each of the three size classes within each DMU and decade by species and palatability class. We averaged values across all plots within each DMU and period. Because these values varied widely and included many zeros, we add one to all values and log transform the averages. This homogenized the residual error variances among taxa and treatments, meeting assumptions of our linear models and facilitating comparisons among species, palatability classes, DMUs, and periods where abundance counts differed greatly. These transformed numbers of small saplings provide a primary indicator for tree regeneration. On this scale, values of 0 reflect sites with no saplings: log(0+1). This was the mode for the more palatable species. We assess demographic inertia between the numbers of small and medium saplings using simple correlations between these transformed values. We also compare the abundance of saplings and larger trees among the palatability classes and decades. This allowed us to evaluate whether saplings numbers in more palatable classes varied independently of tree numbers, as expected if deer control their numbers. We also compare patterns of recruitment among the palatability classes to see whether recruitment in the palatable taxa differs from recruitment in unpalatable *Picea* and *Abies*. We plot and evaluate the distributions of sapling numbers in each species to see whether these differed in shape in a diagnostic way between more and less palatable species (hypothesis b). Note the use of two controls here: the numbers of adult trees in the same taxon and sapling numbers in the unpalatable taxa.

To address question c), we compare the extent to which variation in sapling abundance within palatability classes and species covaries with estimated deer densities within that DMU in the preceding time period. In more browse-sensitive species, we expect the proportion of small saplings to decline in DMUs and decades experiencing higher deer densities but little to no effect of deer density in taxa that deer avoid. Here, we use two complementary approaches. The first used general linear models to evaluate the effects of palatability class and decade on the abundances of small, medium, and large size class trees (questions a and b). This approach implicitly assumes independence in these tree numbers among the DMUs and across the three successive decades of the FIA data. To address the latter assumption, our second approach modeled autocorrelation explicitly using path analysis (see below). We plotted the adjusted means and error bars from the first analysis to compare levels of recruitment among the palatability classes and decades and overall changes in the abundances of small, medium, and larger trees. If deer reduced the densities of saplings in more palatable species, we expect a strong main effect of palatability class on the abundance of small saplings. With regional changes in deer density or another broadly acting ecological factor, we also expect a main effect of decade. Finally, if numbers of small saplings (henceforth “saplings”) in more palatable taxa decline more in decades with more deer, we expect a significant decade by palatability class interaction.

We also tested for deer effects more directly by analyzing variation in small sapling recruitment in relation to both palatability class and previous deer density using DMU estimates from the previous decade as a covariate. Here, again, we expected saplings of more palatable species to be scarcer and for deer abundance to affect sapling abundance more in palatable than in less palatable species (tested using the palatability class x deer interaction term). We also analyzed models for individual species. Because species’ responses to deer varied greatly (deer density x species interaction term F = 8.11, p < 0.001), we analyzed separate models for each taxon. We modeled sapling recruitment as a function of deer density and deer density squared (to account for non-linear effects. We then sequentially dropped non-significant effects in each model to obtain a final model for each species, always retaining the deer effect.

Our final approach analyzed deer effects using path analysis to model tree recruitment in successive decades as a function of sapling numbers and estimated deer densities in the preceding decade (Fig. 7). Path analysis assumes that variables can be placed into causal order, as is the case here given the sequence of events and known effects of herbivory on plants. It also assumes that the modeled predictor variables have linear and additive effects on the dependent variable, assumptions we tested before applying the model. In particular, we summed the (log) number of saplings in palatability class 2 (*Acer rubrum, A. saccharum*, and *Populus tremuloides*) within each decade and DMU and modeled these abundance values as a function of estimated deer densities and sapling numbers in the preceding decade. We applied the model to this palatability class because these species produce abundant seeds and seedlings that provide good indicators for evaluating deer impacts in our region (see below). The path analysis explicitly addresses sequential causal dependencies and the autocorrelation present in these data. We used both R (RStudio: Integrated Development for R, Boston, MA) and JMP (vers 11.2.1) for these analyses (SAS Institute, Cary, NC).

**Table 5.** Results from the path analysis model of recruitment in palatability class 2 (*Acer rubrum* and *saccharum* and *Populus tremuloides*). “Path” refers to the paths labels in Fig. 7 that connect particular predictor and dependent variables. Beta values are the path coefficients (standardized partial regression coefficients). Significance labels for individual predictors as in Table 4. The direct effects of deer on (log) sapling numbers in successive decades are shown in the “Direct effect” column. Indirect effects reflect the delayed effects of deer in an earlier decade as expressed through their effects on later deer abundance and sapling numbers. These involve products of the path coefficients (e.g., the b*f + c*g two-step indirect effects on saplings in 2002 and the e*i + f*j effects on 2011 saplings). Note that total deer effects increase steadily over time as these consistently negative indirect effects have cumulative effects on the number of surviving saplings.

|  |  |  |  |  |  |  |  | Total |
| --- | --- | --- | --- | --- | --- | --- | --- | --- |
|  |  | Dependent |  |  | Direct | Indirect | Indirect (3 | deer |
| Path | Predictor | Var | Beta | Signif | effect | (2 steps) | steps) | effect |
| a | Deer 83 | Saplings 83 | 0.096 | NS |  |  |  |  |
| b | Deer 83 | Deer 96 | 0.639 | *** |  |  |  |  |
| c | Deer 83 | **Saplings 96** | -0.311 | * | -0.311 |  |  | -0.311 |
| d | Saplings 83 | Saplings 96 | 0.447 | *** |  |  |  |  |
| e | Deer 96 | Deer 02 | 0.748 | *** |  |  |  |  |
| f | Deer 96 | **Saplings 02** | -0.270 | * | -0.27 | -0.317 |  | -0.587 |
| g | Saplings 96 | Saplings 02 | 0.465 | *** |  |  |  |  |
| h | Deer 02 | Deer 11 | 0.723 | *** |  |  |  |  |
| i | Deer 02 | **Saplings 11** | -0.417 | ** | -0.417 | -0.429 | -0.337 | -1.183 |
| j | Saplings 02 | Saplings 11 | 0.434 | ** |  |  |  |  |

## 3. Results

### 3.1 Variation in deer densities and sapling abundances

Deer densities varied widely across the region and over time, providing the variance necessary to infer deer effects on sapling numbers (Fig. 2). We also observed substantial variation among sapling numbers across sites within species (Fig. 3). However, these distributions differed in shape among species and palatability groups. The five taxa in palatability classes 1 and 2 showed positive modes and approximately log-Normal distributions (e.g., *Acer rubrum*, Fig. 3a). In contrast, the five species in palatability classes 3 and 4 all showed modes of 0 (reflecting no saplings within the plot) and highly skewed distributions relative to both the less palatable species (Fig. 3b) and to adult trees in the same species. The five more palatable species showed far more skewed distributions of (log) sapling numbers than the less palatable species (means: 1.97 vs. 0.3, Table 1). Sapling numbers were also far more skewed in palatable species relative to adult trees in the same species (mean difference 0.44 in the five less palatable species vs. 1.99 in the more palatable species, paired t-test: t = 3.94, p = 0.003). The presence of adult trees at most sites that lacked palatable saplings suggests that site suitability is not limiting recruitment in these species. We conclude that a syndrome involving very low mean sapling abundance, a mode of 0, and a highly skewed distribution provides a plain and useful signal for inferring strong deer impacts. Surprisingly, this signal emerged for half the taxa studied here. Skew was also noted in seedling counts of some tropical trees and used to predict subsequent demographic dynamics (Feeley et al. 2007).

**Figure 3.**
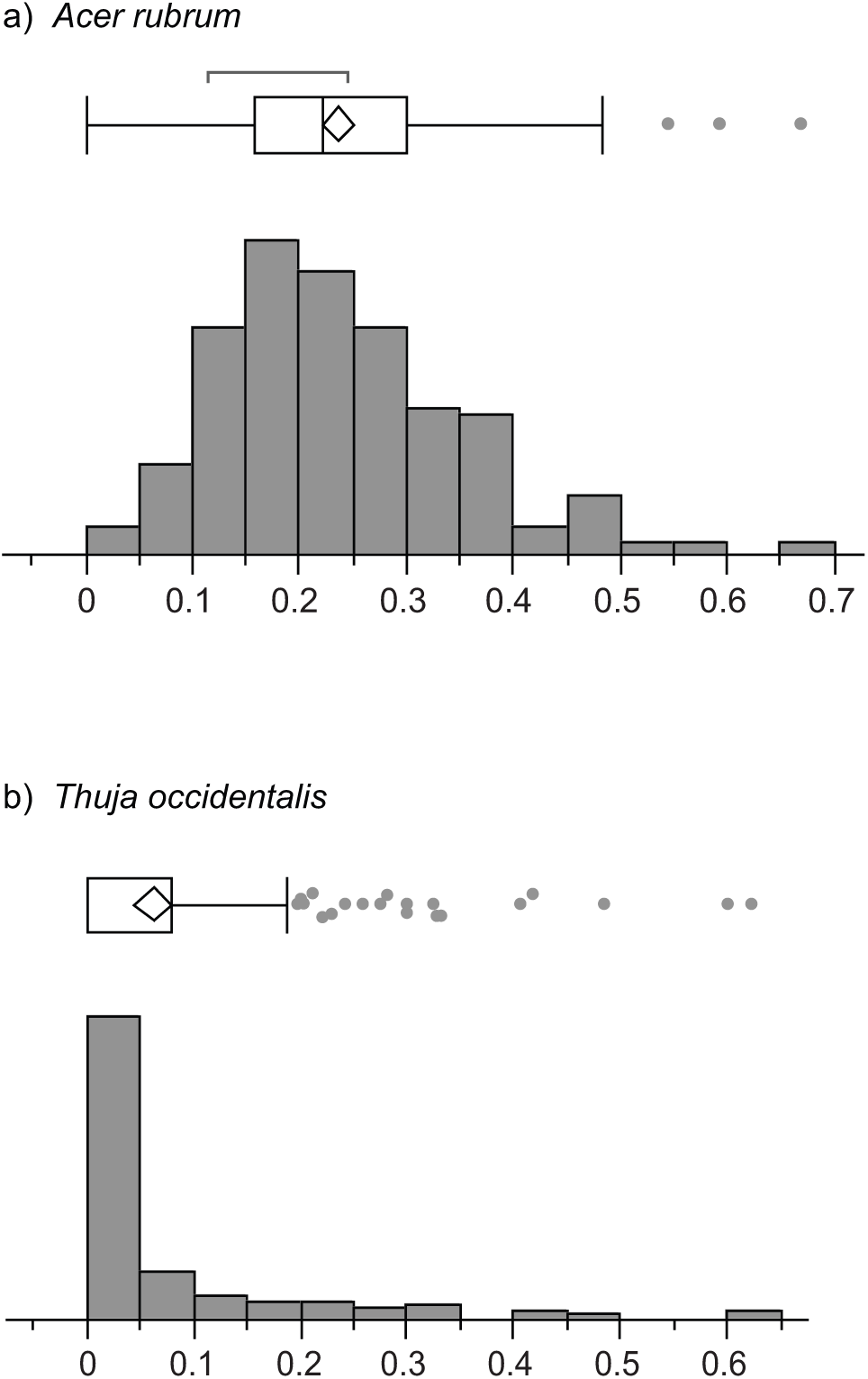
Distributions of red maple and northern white cedar sapling numbers among stands. Frequency histograms show the logarithm of the mean number of small saplings (+1) in all stands. Note the approximately log-Normal distribution of sapling abundances in red maple (a), a prolific seeder of intermediate palatability to deer, and the highly skewed distribution in cedar (b) which is highly palatable and susceptible to deer browse. Mean, variance, skew, and sample sizes for red maple: 0.236, 0.012, 0.816, N=192, and cedar: 0.064, 0.013, 2.52, N = 163.

### 3.2 Palatability classes compared

The abundance of small saplings differed greatly among the four palatability classes and across the four decades (Fig. 4; Table 2). Taxa known to be sensitive to browsing (palatability classes 3 and 4) have far fewer small saplings than taxa that tolerate or repel browsing (classes 1 and 2). The abundances of medium-sized saplings parallel those of the small saplings across decades and the palatability classes (top and middle rows of Fig. 4) where palatability class has a similarly large effect (F=78.2 vs. 66.4, Table 2). We confirmed demographic inertia between size categories using correlations within each species which ranged from +0.51 in *Quercus* to +0.85 in *Abies* (all p < 0.001). In contrast, the abundance of larger trees only loosely reflects palatability. The more palatable classes 3 and 4 had many more adults than juveniles (Fig. 4; Table 2). Thus, the scarcity of saplings in these species does not reflect a lack of seed inputs.

**Figure 4.**
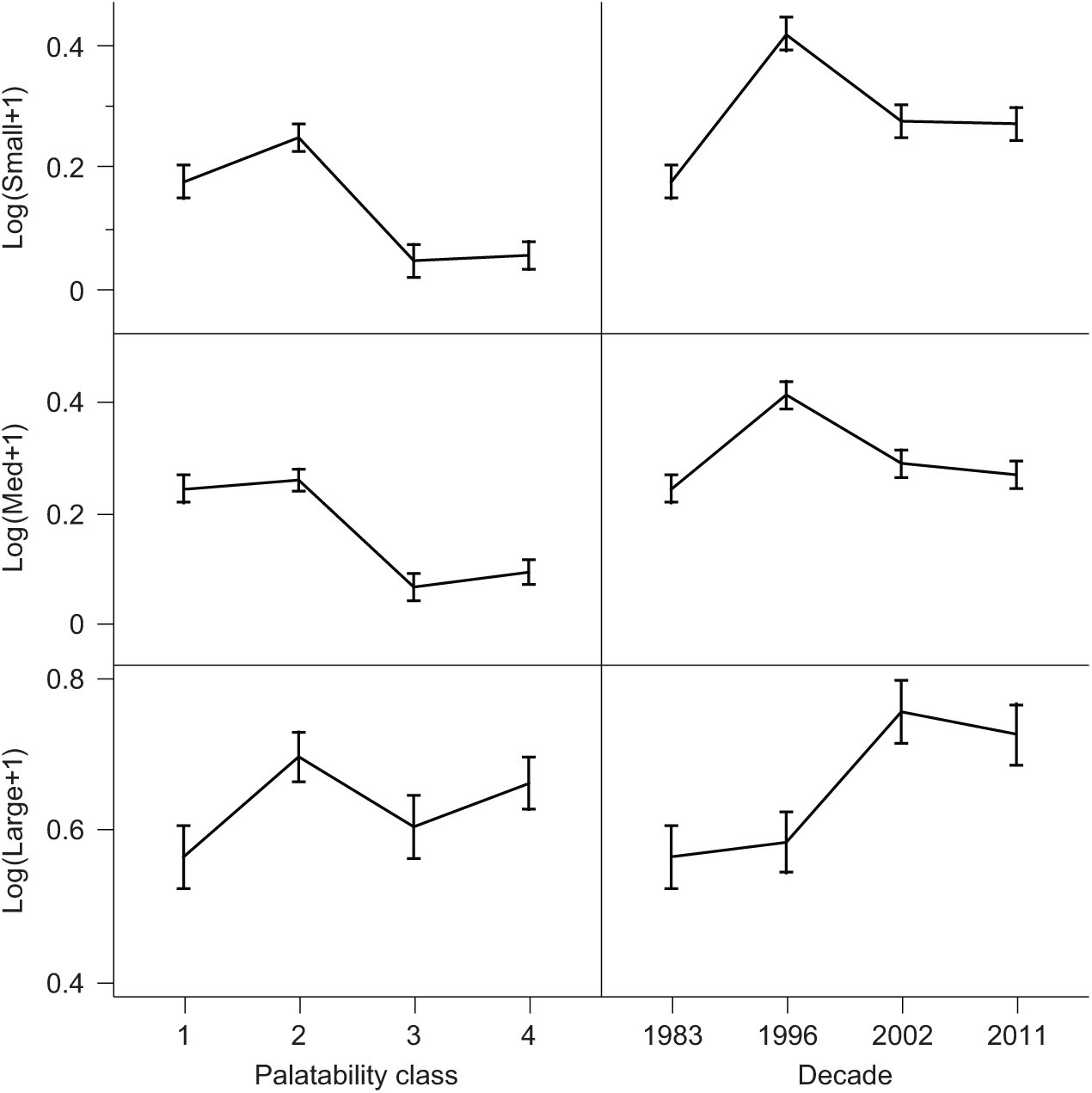
Variation in the numbers of saplings and trees among palatability classes and decades. Values reflect the (log) mean number of trees per stand in all three size classes (small saplings 2.5-5.1cm DBH, medium saplings 5.1-10.2cm DBH, and large sapling >10.2cm DBH). Values are adjusted means correcting for DMU and decade. Error bars show +/- one S.E. Note different scale for the number of large trees.

**Table 2.** Variation in tree abundance among palatability classes. The table shows results from analyses of variance of the logarithm of the number of small, medium, and large sized trees (+1) analyzed over the four palatability classes (1 = least, 4 = most palatable) and decade. The values shown are F-ratios. All values shown are significant at p < 0.001. Adjusted means from these analyses appear in Fig. 3.

| **F-ratios:** |  |  |  |  |
| --- | --- | --- | --- | --- |
| **Factor** | **DF** | **Small** | **Medium** | **Large** |
| Palatability | 3 | 66.38 | 78.16 | 9.81 |
| Decade | 3 | 52.19 | 16.86 | 22.52 |
| Decade*Palatability | 9 | 10.86 | 7.77 | 6.26 |
| **Overall $r^2$ values:** |  | 0.503 | 0.484 | 0.21 |

Across decades, numbers of small saplings increased between 1983 and 1996 and then declined (Fig. 4). This may reflect an initial period of improving regeneration followed by declines in regeneration as increased deer populations (Fig. 2) curtailed regeneration. This apparent lag supports using previous decade deer densities to infer deer effects. Remarkably, these two simple predictors, palatability class and decade, together account for half of the observed variation in sapling abundance (Table 2).

In the models that explicitly include estimated deer density (at the whole DMU level) as a predictor, deer have a strong negative main effect (F = 75.8, p < 0.001) but their effect also varies significantly among the palatability classes (deer x palatability interaction F = 12.5, p < 0.001, Table 3; Fig. 5). Analyses of individual interaction terms show that dramatic declines in sapling numbers as deer increase in palatability class 2 account for most of this effect (Table 3). We expected the relation between sapling numbers and deer density to disappear in unpalatable species but were surprised to also see little response to deer in palatability classes 3 and 4. Again, a model that includes only the effects of DMU average estimated deer density and palatability class accounts for a large proportion (39%) of the total variation observed across sites and decades in small sapling numbers (Table 3). Models with more local estimates of deer abundance would likely do even better.

**Table 3.** Recruitment variation over palatability classes in relation to deer. The table shows results from the general linear model analyzing variation in the (log) number of small saplings in relation to assigned palatability class, estimated deer density during the previous study period, and their interaction. Note that although all four palatability classes differ strongly in small sapling abundance, only class 2 species show significant declines in saplings with increases in estimated deer density. Overall F = 123.3 (df = 7, 1343).

|  | Df | Sum Sq | Mean Sq | F value | Pr(>F) |  |
| --- | --- | --- | --- | --- | --- | --- |
| Palatability | 3 | 429.24 | 143.081 | 249.897 | < 2.20E-16 | *** |
| Deer density | 1 | 43.38 | 43.375 | 75.757 | < 2.20E-16 | *** |
| Deer density:Palatability | 3 | 21.42 | 7.141 | 12.472 | 4.79E-08 | *** |
| Residuals | 1343 | 768.95 | 0.573 |  |  |  |

*Dependent variable : log(small+1)*
|  | Estimate | Std. Error | t value | Pr(>\|t\|) |  |
| --- | --- | --- | --- | --- | --- |
| (Intercept) | 0.450 | 0.032 | 14.124 | < 2.00E-16 | *** |
| Palatability2 | 0.134 | 0.039 | 3.466 | 0.000546 | *** |
| Palatability3 | -0.291 | 0.053 | -5.507 | 4.36E-08 | *** |
| Palatability4 | -0.353 | 0.051 | -6.977 | 4.71E-12 | *** |
| Deer Density | -0.003 | 0.001 | -2.140 | 0.032572 | * |
| Deer density:Palatability2 | -0.007 | 0.002 | -4.071 | 4.95E-05 | *** |
| Deer density:Palatability3 | 0.001 | 0.002 | 0.580 | 0.561692 |  |
| Deer density:Palatability4 | 0.003 | 0.002 | 1.166 | 0.244006 |  |
| Observations | 1350 |  |  |  |  |
| $R^2$ | 0.391 | Adj $R^2$ | 0.388 | | |

**Figure 5.**
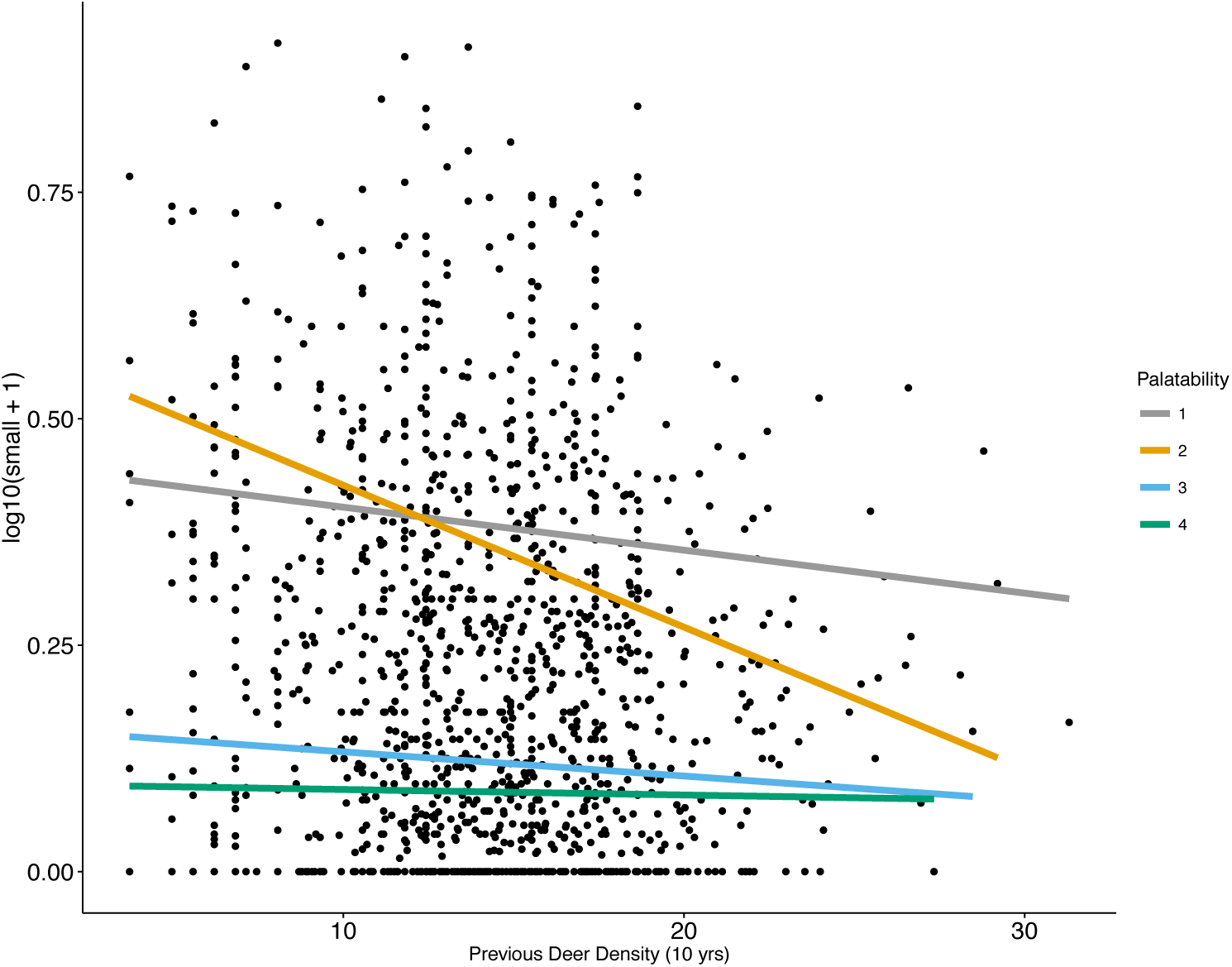
Deer effects on the number of small saplings by palatability class. Deer density estimates are for each DMU in the previous sampling period (about 10 years earlier). Based on the model of sapling numbers in Table 3. The scarcity of small saplings in Palatability classes 3 and 4 precluded significant deer density effects there but class 2 shows a highly significant decline in sapling numbers in relation to deer density (p<0.001).

### 3.3 Species-specific responses

Analyses of individual taxa confirm that species of intermediate palatability show the greatest sensitivity to estimated deer density (Table 4, Fig. 6). Deer effects in *Picea* and *Abies* are negligible, confirming that they serve well as controls. Sapling numbers in five species (mostly in palatability class 2) decline in apparent response to increasing deer density. Sapling recruitment in *Acer rubrum* and *saccharum* and *Populus tremuloides* all declined greatly as deer became more abundant, accounting for the sensitive response to deer in palatability class 2 (Fig. 5). Sapling numbers in red oak (*Quercus rubra*) and yellow birch (*Betula alleghaniensis*) were lower but also declined conspicuously in areas / times of higher deer density. Although we classed balsam fir (*Abies balsamea*) as unpalatable, it showed an almost significant decline in abundance in response to deer (Table 4). We were initially surprised to find no apparent response to deer in *Tsuga canadensis* and *Thuja occidentalis*, two slow-growing conifers known to be highly susceptible to deer browsing (Table 4, Fig. 6). We also found an apparent positive effect of deer on sapling numbers in *Pinus strobus*. These effects, however, were small reflecting the absence of saplings at most sites in these species and corresponding low power for detecting any deer signal. Regeneration was so restricted in these species that intercepts of their fitted regressions (the numbers of small saplings expected when deer are absent) did not differ significantly from zero.

**Table 4.**
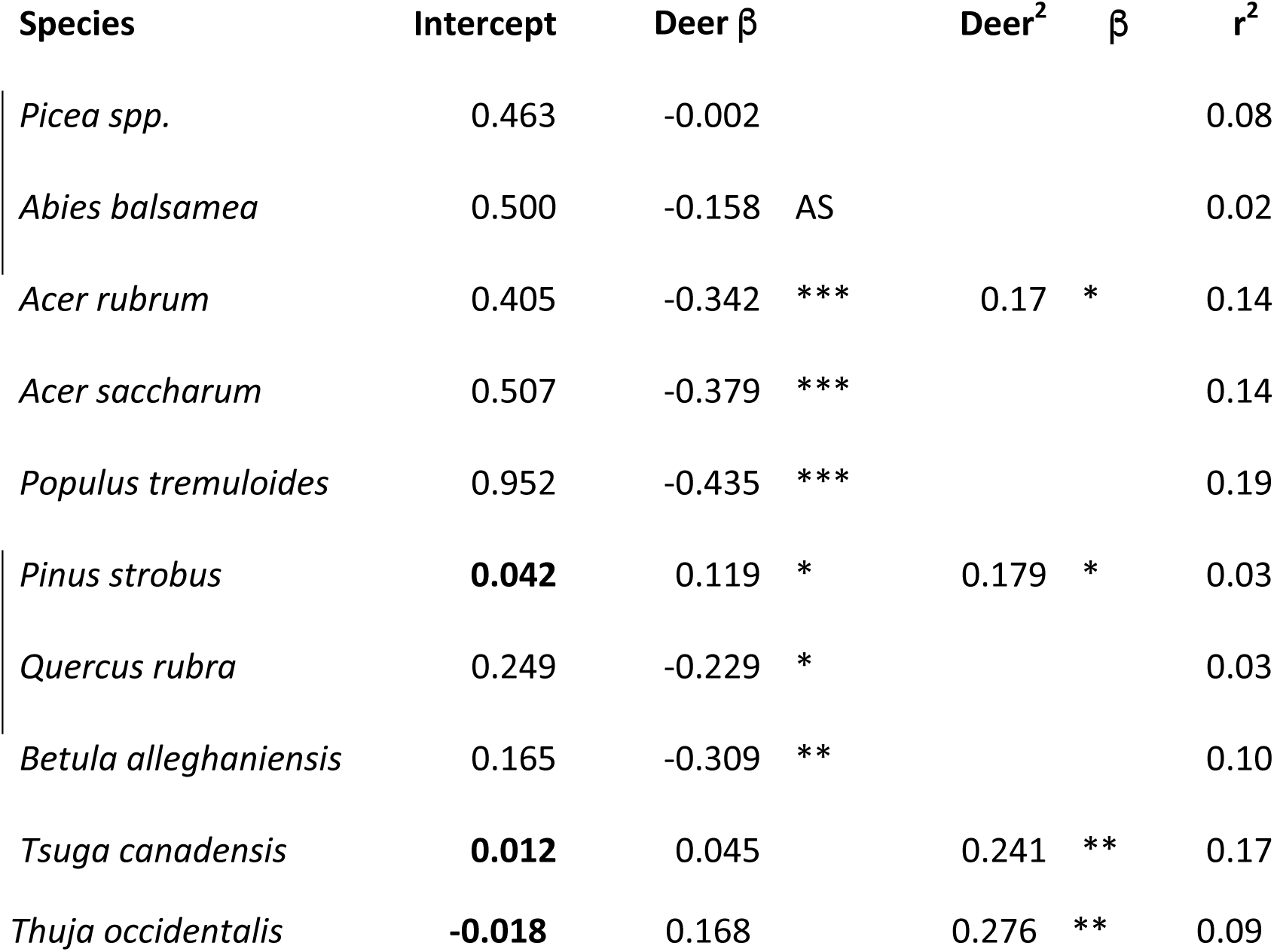
Deer affect sapling recruitment within species. These results derive from the separate models fit to the number of small saplings (log of (number + 1)) for each species (totals within each DMU and time period). Species are listed in order of their palatability classes (Table 1, with classes 1 and 3 marked on left). “Deer β” refers to standardized partial regression coefficients for the linear effects of Wisconsin DNR estimates of deer density within each Deer Management Unit (DMU) during the previous time period (roughly 10 years earlier). Labels show levels of significance: *** p < 0.001, ** p < 0.01, and * p < 0.05, AS p < 0.10. The influence of quadratic (Deer^2^ β) terms are also shown and included in the model when these were significant. Average stand basal area (weighted by each species’ abundance across stands within each DMU) was also included in the model when it was significant (in *Picea*, *Populus*, and *Tsuga*, resulting in beta values of −0.279, −0.14, and 0.471, respectively). The r^2^ terms show overall coefficients of determination for each model. Intercept values show the densities of saplings expected at a deer density (and basal area) of 0 given the model. Values that do not differ significantly from 0 are **bolded**.

**Figure 6.**
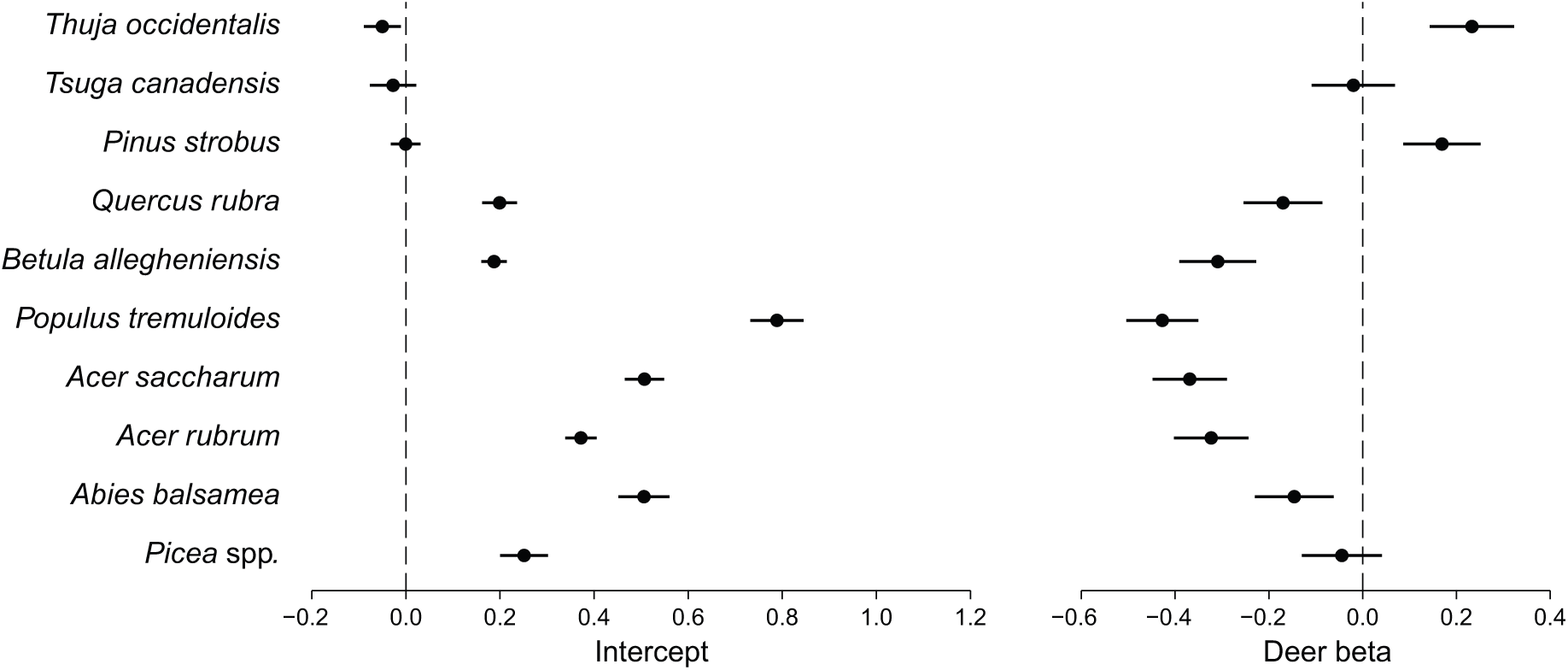
Variation among species in sapling abundance and apparent deer effects. Species appear in order of their predicted palatability to deer from highly palatable conifers (top) to least palatable taxa (*Picea* and *Abies*) at the bottom. The left side shows intercepts from the models fitting (log) small sapling abundance to regional variation in estimated deer densities (from Table 4). Intercepts close to zero (top) reflect species with very few small saplings even at relatively low deer densities. The right side shows the estimated effect of deer (β values) on variation in small sapling numbers by species. Direct deer effects are most easily detected in abundant species of intermediate palatability.

**Figure 7.**
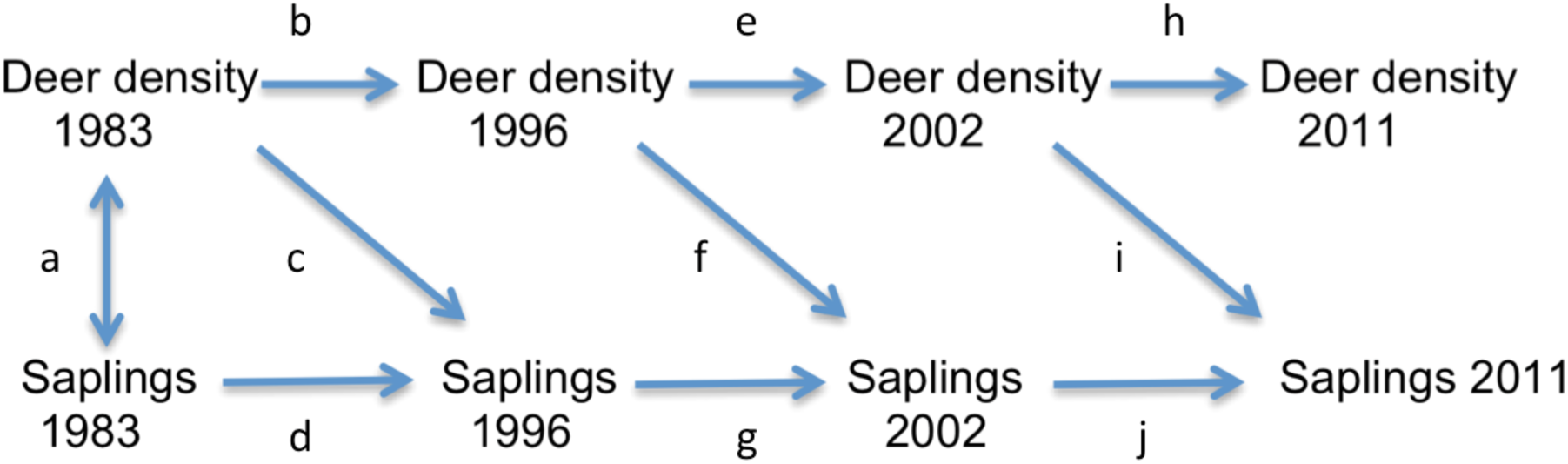
Path diagram showing effects of deer on patterns of sapling recruitment. Tree data reflect sums of the average number of small saplings in palatability class 2 (*Acer rubrum*, *A. saccharum*, and *Populus tremuloides*) across all stands within a particular DMU and decade. Deer densities are Wisc DNR estimates for each DMU and decade. Path coefficients for each lettered path appear in Table 5.

### 3.4 Path analysis

The path analysis (Fig. 7) allowed us to explicitly model sequential time dependencies and the autocorrelation present in our data sets. Deer densities in each decade strongly affected deer densities in the following decade (beta values: 0.64 to 0.75), as did sequential sapling abundances in palatability class 2 (beta = 0.43 to 0.46, Table 5). The direct effects of deer were always appreciable and negative with beta values of −0.27 to −0.42. Interestingly, all the 2- and 3-step indirect effects were also negative and of the same magnitude (betas from −0.32 to −0.43). Thus, over successive decades, as the model includes more information about deer densities and sapling numbers in preceding decades, the indirect effects of deer accumulate to become much larger than the direct effects (Table 5). This strongly implies that the effects of deer analyzed in the preceding GLM models substantially underestimate the actual total effects of deer on tree regeneration in these forests.

## 4. Discussion

The metrics and methods applied in this study all point to the conclusion that browsing by white-tailed deer strongly limits patterns of tree recruitment in the forests of northern Wisconsin. Deer have considerably reduced regeneration in many tree species of intermediate palatability and eliminated it altogether at most sites in several conifer species known to be sensitive to deer browsing. This is highly relevant for deer management in that stands of these conifers provide key habitats for overwintering deer. These stands will likely disappear given this regional failure in regeneration. These effects are not local, temporary, or restricted to a few sensitive species. Rather, they extend across all of northern Wisconsin, cover a 30 year period, and affect most (8 of 10) of the tree species examined. These effects have also begun to modify the composition and structure of the mid-stories of these forests. It will not be simple to limit or reverse these impacts, but without these data and analyses, we could not assess the full scale and extent of the impacts deer are having.

The results of this study plainly indicate that we must use different indicators to detect and monitor deer impacts on forest tree regeneration in different species in this region. Current and previous levels of deer browsing were so heavy in the five most palatable species that small saplings were scarce to absent at most sites. Sapling densities were too low in the highly palatable conifers to accurately measure regeneration or to relate variation in regeneration to deer density. For these species, the absence of small saplings is itself the best indicator, as quantified by very low means, modes of zero, and highly skewed distributions of sapling numbers. Fortunately, these indicators are easily extracted from the FIA data and can be used even when data on deer densities are suspect or lacking.

Species of lower palatability to deer had more abundant saplings allowing us to examine how mean sapling abundance varied with respect to estimated deer density. In these species, we found highly significant negative associations, strongly supporting hypothesis c) and providing an altogether different method for inferring deer impacts. Having these two independent indicators that apply to separate but overlapping sets of species that differ in regeneration success adds range and flexibility to the tools that forest and wildlife managers can use to monitor and assess deer impacts. Together, these metrics support all three hypotheses a), b), and c) that saplings of highly deer-palatable species are now scarce to absent across much of our region and that deer densities strongly affect sapling abundance in several species of lower palatability.

Research in the eastern and Midwestern U.S. has already shown that *Thuja occidentalis*, *Tsuga canadensis*, and *Quercus rubra* are highly palatable to deer and have failed to regenerate in many forests (Strole and Anderson 1992; Boerner and Brinkman 1996; Waller et al. 1996; Alverson and Waller 1997; Rooney et al. 2000, 2002; Collins and Carson 2002). All these studies identify white-tailed deer as the key factor limiting the density or size of seedlings or saplings in these species. The limitation of these studies has only been that they have focused on regeneration at a limited number of sites and documented these failures over limited periods of time. These limitations have allowed some to argue that these failures are local, temporary, and/or only apply to a few highly palatable species. In contrast, our study exploited the vast amount of information contained in the FIA database to show that deer have curtailed sapling recruitment in most of the tree species studied and have had these strong impacts over at least 30 years and a huge region reflecting a diversity of forest types, ownerships, and methods of management. Our results thus complement and amplify results from more local studies by confirming that the common evidence of local regeneration failure they have documented apply much more broadly to multiple species, most sites, and an extended period of time.

Numbers of tree saplings varied strongly among the palatability classes. As predicted, we found many fewer saplings in classes 3 and 4 than in 1 and 2. Decade and palatability class alone account for more than half (51%) of the total variation in regeneration success observed across all tree species studied and all 13,105 sites. For such simple predictors to provide such high explanatory power is rare in complex ecological systems occupying a diverse and heterogeneous region. Substituting estimated deer densities for decade in these models still provided high explanatory power (39%) despite the fact that these DMU estimates reflect spatial averages that ignore all the local variation in deer densities and site conditions affecting tree regeneration within these DMU’s (Wisconsin DNR 2006). As judged by F values, palatability consistently emerged as the most important predictor of sapling abundance and one that always interacted strongly with the deer effect, exactly as predicted. The fact that statistical models that ignore all local variation in soil and light conditions known to greatly affect patterns of tree regeneration had such high explanatory power strongly suggests that deer browsing has emerged as the dominant force controlling patterns of tree sapling recruitment in these forests.

Saplings became progressively scarcer in the higher palatability classes (3 and 4) coincident with the emergence of conspicuous skewed sapling abundance. This high skew with a mode of 0 demonstrates that some factor acted at most sites to eliminate regeneration altogether. Saplings in browse-sensitive *Betula alleghaniensis* and *Quercus rubra* were still abundant enough to show strong deer effects (with betas of −0.31 and −0.23, respectively). Their sapling distributions, too, however, show high skew and an absence of saplings at many sites. Deer exclosures in our region confirm that yellow birch seedlings become scarce where deer are common (Frerker et al. 2013). If saplings in *Betula* and *Quercus* become even more scarce, it will become impossible to detect direct deer effects from among-site variation in sapling abundance.

We are the first to propose using a very low sapling abundance and high skew in (log) sapling numbers relative to adult trees of the same species as plain and simple signatures of deer impacts. We observed that diagnostic signature in half the species studied here, but only in species with higher palatability to deer. We further note that anyone with access to FIA data for their area can easily compute these metrics and that computing these to infer deer impacts requires no data whatsoever on estimated deer densities, local rates of browsing, local seedbed conditions, or any other labor-intensive field measurement. The mode of 0 saplings with very high skew together provide a useful and reliable signature of deer impacts with skew (of log sapling number) emerging as more diagnostic. Feeney et al. (2007) also identified skew as a useful metric as it correctly predicted the direction of population change within two Malasian forests for ~75% of the species. They also found, however, that trends within one forest did not serve to predict demographic changes at the other site. We expect that the far broader sampling enabled by the FIA data should allow more accurate inferences regarding regional trends. This conjecture and the skew and mode metrics should, however, now be tested in other systems.

Trends in our two controls (adults of the same species and the unpalatable taxa) served well to confirm that the metrics we used to infer deer impacts provide robust and reliable signals. The abundances of larger trees were largely uncoupled from patterns of sapling recruitment, confirming that trees are there to provide seed inputs and that sites are indeed suitable for these species. Saplings of unpalatable taxa (*Picea* and *Abies balsamea*) occurred commonly at most sites but their abundance did not covary with deer abundance. These controls support the conjecture that deer rather than some other factor has acted to limit tree recruitment in these forests. Surprisingly, deer may even have acted to depress *Abies* sapling abundance (beta = 0.16, p=0.06). Deer generally only consume *Abies* when other browse is scarce (Dahlberg and Guettinger 1956; Tremblay et al. 2005).

We exploited natural variation in deer density across our region as a “natural experiment” to explore how deer affect patterns of regeneration in our studied species. In our region, densities are lowest in the larger Ojibway and Menominee Indian reservations where densities usually remain less than 4 per km^2^ (Wisc DNR data, R. Rolley). Deer densities clearly also vary greatly in response to local habitat conditions, current and recent hunting pressure, and levels of predation (Leopold 1933). Despite this local variation in deer abundance and the imprecision inherent in the Wis-DNR SAK estimates of deer density, these whole DMU estimates of deer density worked surprisingly well to predict regional variation in the density of saplings in five of the ten taxa we studied (Table 4). These included two important sawtimber species (*Quercus rubra* and *Betula alleghaniensis* in palatability classes 3 and 4, respectively) as well *Acer* and *Populus* with intermediate palatability. The historical depth of the SAK deer estimates and abundance of these species also allowed us to demonstrate both immediate effects of deer on sapling numbers and how deer affected sapling recruitment over successive decades using path analysis. These cumulative impacts acted to multiply total deer impacts in these moderately palatable tree species over the 30 years of this study. Results from the path analysis also suggest that the recent deer effects examined in the simpler models actually underestimate total deer effects by ignoring how deer impacts accumulate over successive decades.

The species showing the strongest responses to deer abundance had intermediate palatability (class 2, *Acer rubrum* and *saccharum* and *Populus tremuloides*). These widespread trees often dominate sites where they occur. They produce prolific numbers of seeds and seedlings that then recruit to become the saplings counted in the FIA surveys. The high numbers of seedlings these species produce could help to swamp local herbivores like deer by improving survival odds for individual seedlings when many are present. Such effects could also cause sapling numbers to decline non-linearly as deer populations increase as many saplings could survive when deer are low relative to seedling numbers but few to none may survive when deer become more abundant. Such effects probably contribute to the high skew we observed in the more palatable species as well as the steep declines in sapling recruitment with deer density in *Acer* and *Populus*. The abundant seedlings and saplings in these species also give us a tool for tracking deer impacts even when deer are dense enough to eliminate regeneration in more palatable species.

Seedlings and saplings of *Pinus strobus*, *Tsuga canadensis* and *Thuja occidentalis* are all highly vulnerable to deer that often browse preferentially on these conifers, particularly in winter (Dahlberg and Guttinger 1956; Alverson and Waller 1997; Rooney et al. 2000; Cornett et al. 2000; Rooney et al. 2002; Townsend et al. 2002; Vila et al. 2003). Saplings of these species were so scarce across the region that no saplings occurred at most sites. Their scarcity prevented us from being able to detect any decline in sapling numbers as deer densities increased. We even observed weak positive relationships in two cases (though the intercepts in these cases were indistinguishable from zero). It would be useful in this context to do numerical simulations to reduce (“rarify”) sapling data to see when clear negative relationships drop to the point of non-significance.

We did not examine the effects of management practices in this study but know that management can enhance seedling establishment in many of the palatable species, e.g., by providing substrates suitable for germination and early growth (Rooney et al. 2002, Curtis 1959, Doepker and Ozoga 1990). Improving seedling establishment, however, does little to improve recruitment to size classes above 2m in height in the shade-tolerant evergreens we studied as slow-growing seedlings must pass through a long period of vulnerability to deer before they escape the “molar zone” (Waller et al. 1996; Rooney et al. 1998, 2000). Silvicultural treatments that enhance light conditions, however, might accelerate seedling growth enough to improve recruitment in some cases‥

Accepting both the scarcity and skew of sapling numbers and negative effects in the linear models as signals of deer impacts, we find strong evidence for these in eight of the ten taxa we studied. Populations of slow growing tree species take 70+ years to recover from prolonged browsing (Frelich and Lorimer 1985; Anderson and Katz 1993). Thus, deer impacts manifest now will persist for many decades to a century or more (McGarvey et al. 2013). Once saplings have failed to regenerate for several decades, the abundance of those species in the canopy will decline. This, in turn, will reduce the input of seeds, further depressing seedling recruitment. The slow rate of these processes delays both our ability to recognize that deer impacts are occurring and our ability to reverse such declines already underway. Thus, we should be vigilant in monitoring patterns of regeneration and addressing causes of regeneration failure if we intend to sustain the diversity and relative abundance of trees that in these forests and the commercial value they represent.

Tree regeneration across northern Wisconsin is now somewhat limited by deer browsing in three of the ten taxa examined, strongly limited in another two, and severely limited in three conifer species. Different signatures of deer browsing are evident in these species depending on seedling abundance, how susceptible their seedlings are to browsing and their rates of growth. These metrics should now be tested in other systems to confirm that FIA data can be put to similar use elsewhere. Because FIA data are already systematically collected for other purposes, using them to estimate deer impacts brings high power at low cost. We gain additional power when we also have access to reliable data on deer densities extending over several decades.

The absence of any routine widespread monitoring system has stymied our ability to reliably infer the extent and severity of deer impacts on forest ecosystems. State and federal agencies can now capitalize on the FIA data to routinely monitor deer impacts in a systematic and sustained way. Sharing these results widely with foresters, hunters, and the public could improve public understanding of deer threats and public support for professional management actions to address those threats. The results presented here strongly support the need to limit deer densities to promote healthy forest regeneration. As always, we should check assumptions, use controls, and accept the limitations involved in using FIA and DNR data. Foresters, forestland owners, and wildlife agency personnel may be keen to use these results to enhance efforts to control deer populations. Limited “band-aid” solutions to browsing such as fencing deer out or protecting individual seedlings are expensive and do not serve to protect other elements of plant and animal diversity sensitive to deer impacts. Once we better understand how forests and wildlife populations interact, we should be able to better manage both as an integrated and dynamic system.

## Acknowledgements

We thank the U.S. Forest Service FIA program and particularly C. Barnett and E. LaPoint for their assistance in accessing and using the FIA data. R. Rolley of the WI DNR helped us to access and interpret the deer population data. T. Van Deelen helped us understand the utility and limitations of the SAK deer estimates. J. Ash and D. Li provided statistical advice. R. Toczydowski, A. Sabo, and G. Sonnier provided helpful comments on the manuscript. S. Friedrich redrew the figures. DMW is grateful for the favorable working conditions and sabbatical support provided by O. Ronce, I. Olivieri, the LABex program, and the ISEM group at the Université de Montpellier, France.

## Graphical Abstract

White-tailed deer (*Odocoileus virginianus*) had substantial cumulative impacts on patterns of forest tree recruitment across most of northern Wisconsin between 1983 and 2013. The diagram shows the path model used to trace direct and cumulative effects of deer on patterns of sapling recruitment in *Acer rubrum*, *Acer saccharum*, and *Populus tremuloides* over this interval. Deer densities refer to 10-year average estimates from Wisconsin DNR’s Sex-Age-Kill model as customized for each of 48 Deer Management Units in northern Wisconsin. Saplings refers to the logged sums of 2.5–5cm DBH sapling numbers in these species as tallied over matching 10-year intervals from U.S. Forest Service FIA survey data on tree numbers and growth within 13,105 plots in this region. See Figure 7 and Table 5. Image of deer used with permission by the Integration and Application Network, Univ. of Maryland Center for Environmental Science (ian.umces.edu/imagelibrary/).

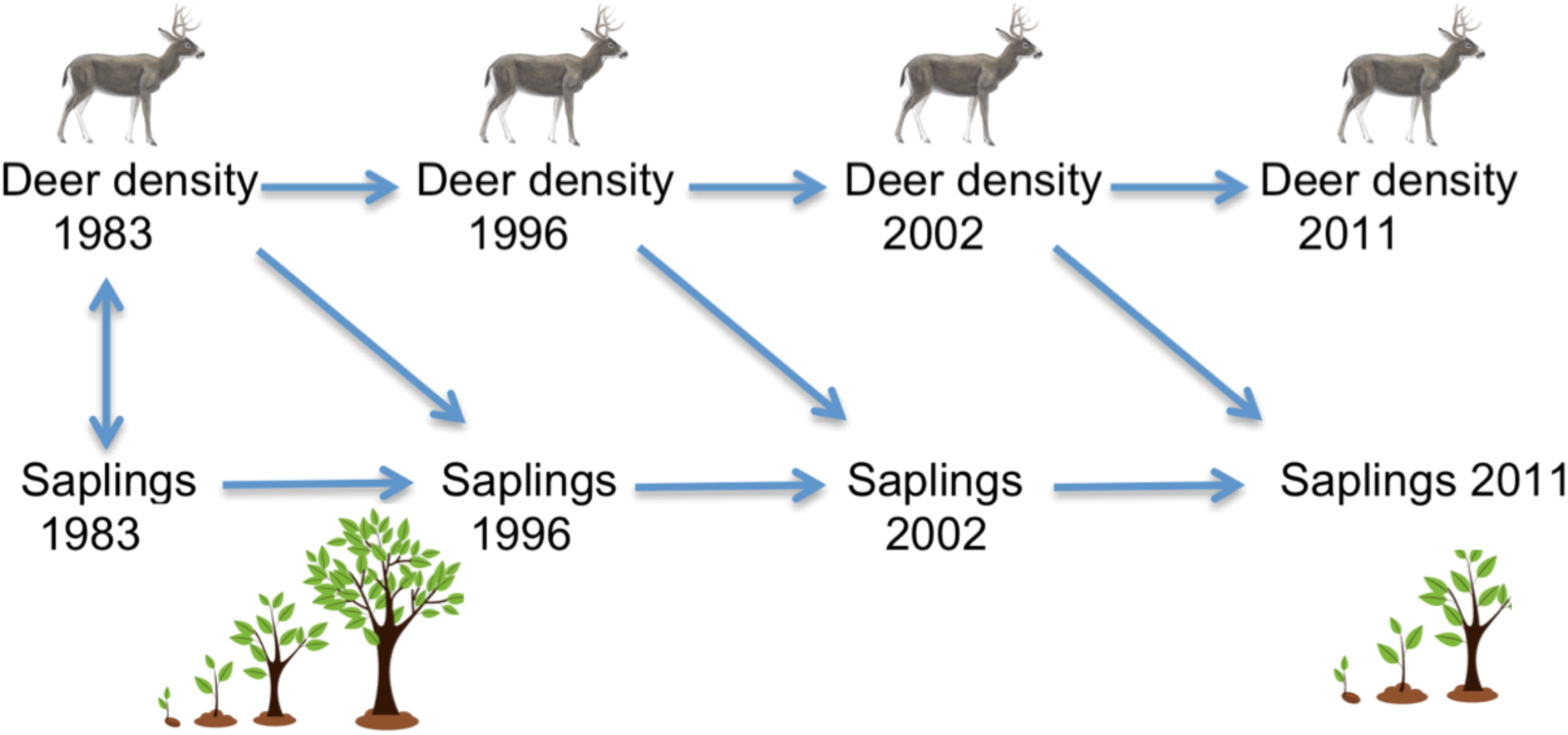

## REFERENCES CITED

Aarssen, L. W., & Epp, G. 1990. Neighbor manipulations in natural vegetation - a review. J Veg Sci, 1: 13–30. doi:10.2307/3236049

Allombert, S., Gaston, A.J., and Martin, J.-L. 2005. A natural experiment on the impact of overabundant deer on songbird populations. Biol. Conserv. 126: 1–13.

Alverson, W.S., and Waller, D.M. 1997. Deer populations and the widespread failure of hemlock regeneration in northern forests. *In* The science of overabundance: deer ecology and population management. Edited by W. McShea and J. Rappole. Smithsonian Inst. Press, Washington, D.C. pp. 280–297.

Anderson, R.C., and Katz, A.J. 1993. Recovery of browse-sensitive tree species following release from white-tailed deer (*Odocoileus virginianus* Zimmerman) browsing pressure. Biological Conservation 63(3): 203–208.

Anderson, R.C., and Loucks, O.L. 1979. White-tail deer (*Odocoileus virginianus*) influence on the structure and composition of *Tsuga canadensis* forests. J. Appl. Ecol. 16: 855–861.

Balgooyen, C.P., and Waller, D.M. 1995. The use of *Clintonia borealis* and other indicators to gauge impacts of White-tailed Deer on plant communities in northern Wisconsin, USA. Natural Areas Journal 15: 308–318.

Berteaux, D., Crête, M., Huot, J., Maltais, J., and Ouellet, J.-P. 1998. Food choice by white-tailed deer in relation to protein and energy content of the diet: a field experiment. Oecologia 115: 84–92. doi: 10.1007/s004420050494.

Boerner, R.E.J., and Brinkman, J.A. 1996. Ten years of tree seedling establishment and mortality in an Ohio deciduous forest complex. Bull. Torrey Bot. Club 123: 309–317.

Bratton, S.P. 1979. Impacts of white-tailed deer on the vegetation of Cades Cove, Great Smoky Mountains National Park. Proc. Ann. Conf. Southeastern Assoc. of Fish & Wildlife Agencies 33: 305–312.

Bressette, J.W., Beck, H., and Beauchamp, V.B. 2012. Beyond the browse line: complex cascade effects mediated by white-tailed deer. Oikos. doi: 10.1111/j.1600-0706.2011.20305.x.

Cardinal, E., Martin, J.L., and Côté‥, S.D. 2012. Large herbivore effects on songbirds in boreal forests: lessons from deer introduction on Anticosti Island. Ecoscience 19: 38–47.

Collins, R.J., and Carson, W.P. 2002. The fire and oak hypothesis: incorporating the effects of deer browsing and canopy gaps. *In* 13th Central Hardwood Forest Conference. Edited by J.W. Van Sambeek and J.O. Dawson and F.J. Ponder and E.F. Loewenstein and J.S. Fralish. USDA, Forest Service, North Central Research Station, St. Paul, MN, Urbana, IL. p. 565.

Côté, S.D., Rooney, T.P., Tremblay, J.-P., Dussault, C., and Waller, D.M. 2004. Ecological impacts of deer overabundance. Ann. Rev. Ecol. Evol. System. 35: 113–147.

Curtis, J. 1959. The Vegetation of Wisconsin. The University of Wisconsin Press, Madison, Wisconsin.

Dahlberg, B. L. and Guettinger, R. C. 1956. Winter Range Condition Surveys. In The White-Tailed Deer in Wisconsin (pp. 156–180).

Dalling, J. W., Davis, A. S., Schutte, B. J., & Elizabeth Arnold, A. 2011. Seed survival in soil: interacting effects of predation, dormancy and the soil microbial community. Journal of Ecology 99: 89–95.

deCalesta, D.S. 1994. Effect of white-tailed deer on songbirds within managed forests in Pennsylvania. J. Wildlife Mgmt. 58: 711–717.

Diamond, J.M. 1983. Laboratory, field, and natural experiments. Nature 304: 586–587.

Doepker, R.V., Ozoga, J.J. 1990. Wildlife values of northern white cedar. In: Lantagne, D.O. (Ed.), Proceedings of the Northern White Cedar in Michigan Workshop, Sault St. Marie, MI, pp.

Feeley, K. J., Davies, S. J., Noor, M. N. S., Kassim, A. R., & Tan, S. (2007). Do current stem size distributions predict future population changes? An empirical test of intraspecific patterns in tropical trees at two spatial scales. Journal of Tropical Ecology. 23: 191. http://doi.org/10.1017/S0266467406003919.

FIA DataMart. <http://www.fia.fs.fed.us/tools-data/>. Visited March 2015.

Frelich, L.E., Lorimer, C.G., 1985. Current and predicted long-term effects of deer browsing in hemlock forests in Michigan, U.S.A. Biol. Conserv. 34, 99–120.

Frerker, K.L., Sabo, A., and Waller, D.M. 2014. Long-term regional shifts in plant community composition are largely explained by local deer impact experiments. PLoS-ONE 9(12): e115843. doi: 10.1371/journal.pone.0115843.

Fuller, R.J. 2001. Responses of woodland birds to increasing numbers of deer: a review of evidence and mechanisms. Forestry 74: 289–298.

Gill, R.M.A., and Beardall, V. 2001. The impact of deer on woodlands: the effects of browsing and seed dispersal on vegetation structure and composition. Forestry 74: 209–218.

Healy, W.M. 1971. Forage preferences of tame deer in a northwest Pennsylvania clearcutting. Journal of Wildlife Management 35: 717–723.

Hobbs, N.T. 1996. Modification of ecosystems by ungulates. J. Wildl. Manage. 60: 695–713.

Kolb, A., Ehrlen, J. & Eriksson, O. 2007. Ecological and evolutionary consequences of spatial and temporal variation in pre-dispersal seed predation. Perspectives in Plant Ecology Evolution and Systematics 9: 79–100.

Leopold, A. 1933. Game Management. University of Wisconsin Press.

Leopold, A., Sowls, K., and Spencer, D.L. 1947. A survey of over-populated deer ranges in the United States. Journal of Wildlife Management 11: 162–177.

Lorenzetti, F., Delagrange, S., Bouffard, D., & Nolet, P. 2008. Establishment, survivorship, and growth of yellow birch seedlings after site preparation treatments in large gaps. Forest Ecology and Management 254: 350–361.

Marquis, D.A. 1981. Effect of deer browsing on timber in Allegheny hardwood forests of northwestern Pennsylvania. USDA Forest Service Research Report. NE-47.

Martin, J.-L., Stockton, S.A., Allombert, S., and Gaston, A.J. 2012. Top-down and bottom-up consequences of unchecked ungulate browsing on plant and animal diversity in temperate forests: lessons from a deer introduction. Biological Invasions doi: 10.1007/s10530-009-9628-8.

McGarvey, J.C., Bourg, N.A., Thompson, J.R., McShea, W.J., and Shen, X. 2013. Effects of Twenty Years of Deer Exclusion on Woody Vegetation at Three Life-History Stages in a Mid-Atlantic Temperate Deciduous Forest. Northeastern Naturalist 20: 451–468.

McShea, W.J., and Rappole, J.H. 2000. Managing the abundance and diversity of breeding bird populations through manipulation of deer populations. Conservation Biology 14: 1161–1170.

Millspaugh, J. J., Skalski, J. R., Townsend, R. L., Diefenbach, D. R., Boyce, M. S., Hansen, L. P., and Kammermeyer, K. 2009. An Evaluation of Sex-Age-Kill (SAK) Model Performance. Journal of Wildlife Management, 73: 442–451.

Mladenoff, D.J., and Stearns, F. 1993. Eastern Hemlock regeneration and deer browsing in the northern Great Lakes region: A re-examination and model simulation. Cons. Biol. 7: 889–900.

Norton, A.S., Diefenbach2, D.R., Rosenberry, C.S., Wallingford, B.D., 2013. Incorporating harvest rates into the sex-age-kill model for white-tailed deer. Journal of Wildlife Management 77, 606–615.

Nuttle, T., Ristau, T. E., and Royo, A. 2014. Long-term biological legacies of herbivore density in a landscape-scale experiment: Forest understoreys reflect past deer density treatments for at least 20-years. Journal of Ecology 102: 221–228.

Ostfeld, R.S., Jones, C.G., and Wolff, J.O. 1996. Of mice and mast: ecological connections in eastern deciduous forests. BioScience 46: 323–330.

Persson, I. L., Danell, K., & Bergström, R. 2000. Disturbance by large herbivores in boreal forests with special reference to moose. In Annales Zoologici Fennici, pp. 251–263. Finnish Zoological and Botanical Publishing Board.

Rooney, T.P. 2001. Impacts of white-tailed deer to forest ecosystems: a North American perspective. Forestry 74: 201–208.

Rooney, T.P., McCormick, R.J., Solheim, S.L., and Waller, D.M. 2000. Regional variation in recruitment of eastern hemlock seedlings in the Southern Superior Uplands Section of the Laurentian Mixed Forest Province, USA. Ecological Applications 10: 1119–1132.

Rooney, T.P., Solheim, S.L., and Waller, D.M. 2002. Factors influencing the regeneration of northern white cedar in lowland forests of the Upper Great Lakes region, USA. Forest Ecology and Management. 163: 119–130.

Rooney, T.P., and Waller, D.M. 1998. Local and regional variation in hemlock seedling establishment in forests of the upper Great Lakes region, USA. Forest Ecology and Management 111: 211–224.

Rooney, T.P., and Waller, D.M. 2003. Direct and indirect effects of deer in forest ecosystems. Forest Ecology & Management 181: 165–176.

Russell, F. L., Zippin, D. B. and N. L. Fowler. 2001. Effects of white-tailed deer (Odocoileus virginianus) on plants, plant populations and communities: a review. American Midland Naturalist. 146: 1–26.

Stormer, F.A., and Bauer, W.A. 1980. Summer forage use by tame deer in northern Michigan. J. Wildlife Management 44: 98–106.

Strole, T. A., and Anderson, R.C. 1992. White-tailed deer browsing: species preferences and implications for central Illinois forests. Nat. Areas J. 12:139–144.

Townsend, D. S., Seva, J.S., Hee-Seagle, C., and Mayers, G. 2002. Structure and composition of a northern hardwood forest exhibiting regeneration failure. Bartonia 61:1–13.

Tremblay, J.-P., Thibault, I., Dussault, C., Huot, J., and Côté, S.D. 2005. Long-term decline in white-tailed deer browse supply: can lichens and litterfall act as alternative food sources that preclude density-dependent feedbacks. Canadian Journal of Zoology 83: 1087–1096.

USDA Forest Service. 2014. Forest Products. <http://www.fs.fed.us/research/forest-products/> Visited November 2015.

Vila, B., Torre, F., Martin, J.L., and Guibal, F. 2003. Response of young *Tsuga heterophylla* to deer browsing: developing tools to assess deer impact on forest dynamics. Trees-Structure and Function 17: 547–553.

Waller, D.M., Alverson, W.S., and Solheim, S. 1996. Local and regional factors influencing the regeneration of eastern hemlock. *In* Regional conference on ecology and management of eastern hemlock. Edited by G. Mroz and J. Martin. Michigan Technological University, Iron Mountain, MI. pp. 73–90.

Waller, D.M., and Alverson, W.S. 1997. The white-tailed deer: a keystone herbivore. Wildlife Society Bulletin 25: 217–226.

Waller, D.M. 2013. Effects of deer on forest herb layers. *In* The Herbaceous Layer in Forests of Eastern North America, 2nd ed. Edited by F.S. Gilliam and M.R. Roberts. Oxford University Press, New York.

White, M.A., 2012. Long-term effects of deer browsing: Composition, structure and productivity in a northeastern Minnesota old-growth forest. Forest Ecology & Management 269, 222–228.

Wisconsin DNR (2006). SAK audit report.

Wisconsin DNR (2015a). Deer abundance and densities in Wisconsin deer management units. <http://dnr.wi.gov/topic/hunt/maps.html>. Visited Aug 2015.

